# A Cytoskeletal Protein Complex is Essential for Division of Intracellular Amastigotes of *Leishmania mexicana*

**DOI:** 10.1101/2020.04.29.068445

**Authors:** Felice D. Kelly, Khoa D. Tran, Jess Hatfield, Kat Schmidt, Marco A. Sanchez, Scott M. Landfear

## Abstract

Previous studies in *Leishmania mexicana* have identified the cytoskeletal protein KHARON as being important for both flagellar trafficking of the glucose transporter GT1 and for successful cytokinesis and survival of infectious amastigote forms inside mammalian macrophages. KHARON is located in three distinct regions of the cytoskeleton: the base of the flagellum, the subpellicular microtubules, and the mitotic spindle. To deconvolve the different functions for KHARON, we have identified two partner proteins, KHAP1 and KHAP2, that associate with KHARON. KHAP1 is located only in the subpellicular microtubules, while KHAP2 is located at the subpellicular microtubules and the base of the flagellum. Both the *KHAP1* and *KHAP2* null mutants are unable to execute cytokinesis but are able to traffic GT1 to the flagellum. These results confirm that KHARON assembles into distinct functional complexes and that the subpellicular complex is essential for cytokinesis and viability of disease-causing amastigotes but not for flagellar membrane trafficking.

## Introduction

*Leishmania* are parasitic protists responsible for an estimated 12 million infections worldwide with pathologies ranging from self-healing cutaneous disease to fatal visceral leishmaniasis (1). These parasites have various developmentally distinct life cycle stages (2), but two major forms amenable to laboratory investigation are the promastigotes that inhabit the midgut of the sand fly vector and the amastigotes that live inside parasitophorous vacuoles within mammalian host macrophages and cause disease. Promastigotes are elongated, spindle-shaped cells of ~15 µm length with a single extended flagellum, and they are highly motile. In nature, infectious metacyclic form promastigotes are delivered to vertebrate hosts by a bite of the sand fly vector. They are subsequently taken up by macrophages and targeted to acidic phagolysosomal vesicles where they transform into oval shaped amastigotes of ~1 - 5 µm that are nonmotile and have a short flagellum that barely extends from the anterior end of the cell body.

Previous studies from our laboratory focusing on membrane transport proteins that mediate uptake of important nutrients in *Leishmania mexicana* parasites identified a unique glucose/hexose transporter isoform that traffics selectively to the promastigote flagellar membrane (3) and was designated GT1. Subsequent investigations on how this permease is selectively targeted to the flagellar membrane identified both a sequence within the N-terminal hydrophilic domain of GT1 that is required for flagellar targeting (3) and a novel protein, named KHARON, or KH (4), that interacts with this targeting sequence and is required for efficient targeting of GT1 to the flagellar membrane. The null mutant generated when the *KHARON* or *KH* gene was deleted, named ∆*kharon* or hereafter ∆*kh*, was strongly impaired in trafficking of GT1 to the flagellum but was without other notable phenotypes in the promastigote stage. Strikingly, the ∆*kh* null mutants were able to infect host macrophages as well as wild type promastigotes, but the intracellular amastigotes died over the course of 7 days post-infection (4), and ∆*kh* null mutants were avirulent following infection of BALB/c mice (5). The intracellular amastigotes were able to replicate nuclei, but parasites could not undergo cytokinesis, leading to multinucleate, multiflagellated amastigotes that eventually disintegrated and were thus unable to support a productive infection (5). Localization studies indicated that KH is a cytoskeletal-associated protein and is attached to i) microtubule-based structures at the base of the flagellum (4), ii) the network of subpellicular microtubules that subtend the cytosolic side of the plasma membrane throughout the parasite cell body (4), and iii) mitotic spindles (6).

The three subcellular loci for KH raise the question of how localization of this protein may be associated with its different functions. Thus, KH located at the base of the flagellum could participate in trafficking of GT1 from the flagellar pocket membrane (7), where membrane proteins are first trafficked during biosynthesis, into the contiguous flagellar membrane. KH may promote passage of GT1 through a ‘periciliary diffusion barrier’ that prevents mixing of flagellar membrane proteins with other proteins from the surface membrane and has been functionally observed in multiple ciliated eukaryotes (8). However, since GT1 is not stably expressed in intracellular amastigotes (9), trafficking of this protein to the amastigote flagellum is not likely to be responsible for the fatal phenotype of ∆*kh* null mutants inside macrophages. While KH could be responsible for trafficking of other unknown proteins into the amastigote flagellum, a distinct possibility is that the failure of ∆*kh* amastigotes to undergo cytokinesis may reflect a function of KH located at other sites within the parasite, either on the subpellicular cytoskeleton or in the mitotic spindle. Furthermore, KH might be associated in different multiprotein complexes with distinct subunits and unique properties at each of its three subcellular loci. If this hypothesis is correct, it may be possible to identify unique KH partners that could help deconvolve the multiple functions of this protein and ascribe different functional roles to distinct KH complexes. As a working hypothesis, we suggest that there may be three such complexes, a KH Complex 1 located at the base of the flagellum, a KH Complex 2 located on the subpellicular microtubules, and a KH Complex 3 associated with the mitotic spindle.

To test the multiple complex hypothesis, we initiated a search for molecular partners of KH. The initial strategy employed BioID (10), a method in which the mutant biotin ligase, BirA*, was fused to the N-terminus of KH to biotinylate other proteins that are in close proximity, including potential molecular partners. The other approach was to tag KH with a tandem affinity purification (TAP) tag and identify other proteins that co-purify with the tagged fusion protein (11). This approach has led to the identification of two ‘KH associated proteins’, KHAP1 and KHAP2, that are molecular partners of KH and co-localize with KH at the subpellicular microtubules but not at the mitotic spindle and, for at least one of them, not at the base of the flagellum. Gene knockouts of *KHAP1* or *KHAP2* block cytokinesis of amastigotes but do not affect trafficking of GT1 to the promastigote flagellar membrane, establishing the role of a unique KH complex at the subpellicular microtubules in cytokinesis of the disease-causing stage of the parasite life cycle.

## Results

### Identification of KH-associated proteins

To identify molecular partners of KH, we first employed the proximity-dependent labeling technique BioID to identify proteins that are found near KH in live cells (10). This method is based on the observation that the promiscuous biotin ligase BirA* biotinylates other proteins within a ~10 nm radius of a relevant BirA*-fusion protein (12). When *L. mexicana* promastigotes expressing BirA*::KH were incubated with biotin, several proteins were biotinylated, as demonstrated by the Western blot in Fig. 1A. This result supports the notion that KH associates with multiple other proteins. We used streptavidin-agarose resin to isolate biotinylated proteins from wild-type and BirA*::KH-expressing parasites, and identified the streptavidin-bound proteins by tandem mass spectrometry. The proteins most enriched in the BirA*::KH sample are listed in Fig. 1B. These data represent two replicate experiments in which each of these proteins was enriched by at least 10 counts in the BirA*::KH compared to wild type sample in mass spectrometry. The relative rank of each identified protein in each of the replicate experiments is listed on the right. Detailed ranking information can be found in Table S1.

**Figure 1.**
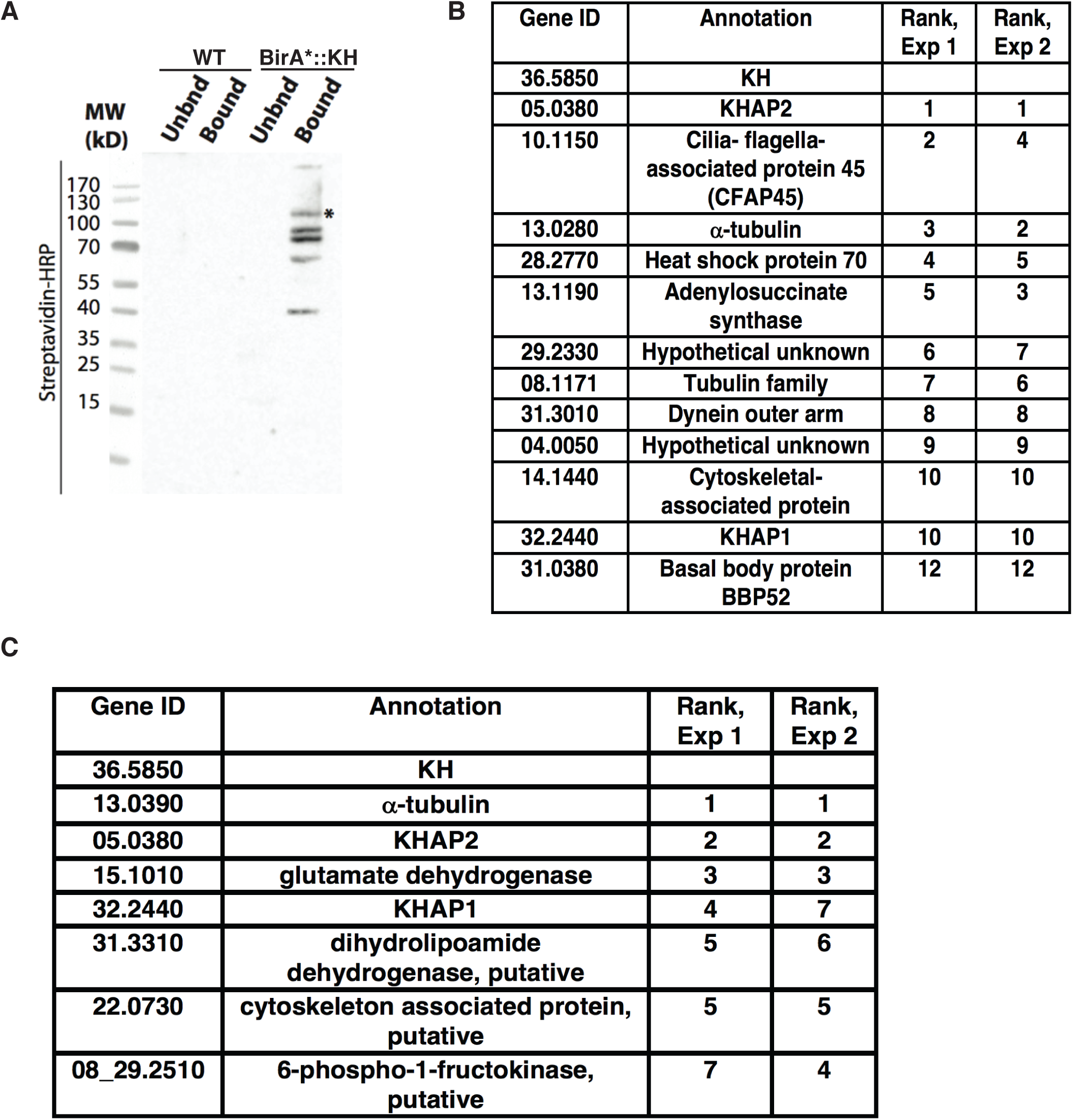
Identification of candidate KH-interacting proteins. *A*, Proximity biotinylation with BirA*-tagged KH. Both wild type *L. mexicana* promastigotes (*W*T, not expressing BirA*) and promastigotes expressing BirA*::KH (*BirA*-KH*) were incubated with biotin, and lysates were subjected to affinity chromatography and eluted from streptavidin-agarose resin. Equivalent fractions of both resin-unbound (*Unbnd*) and bound and eluted (*Bound*) material were separated by SDS-PAGE, blotted onto a nitrocellulose membrane, and probed with streptavidin-horseradish peroxidase (*Streptavidin-HRP*). The *band represents the BirA*::KH bait protein. *B*, BioID hits for BirA*::KH. Two experiments were performed, and spectral counts from wild type samples were subtracted from counts from BirA*::KH-expressing promastigotes. Proteins that showed the largest differential in spectral counts in each of the two experiments are listed, with their relative rank in each experiment indicated. For simplicity, gene IDs are listed without the prefix LmxM used in the TriTrypDB genome database. Proteins with a difference larger than 10 spectral counts in both experiments are included in this list, with the addition of the protein corresponding to gene ID 31.0380, listed despite the differential count value of <10 for Experiment 2, due to its potential interest as a likely basal body protein. More detailed experimental results are reported in Table S1 of Supporting Information. *C*, Tandem-affinity purification of hexa-histidine-biotinylation domain-hexa-histidine-tagged KH (HBH::KH), followed by tandem mass tagging and tandem mass spectrometry identification. Results of two replicate experiments comparing parasites expressing or not expressing HBH::KH are shown, and the proteins with the greatest differential in tandem-mass-tag intensities common to both replicates listed as in (B). This analysis excluded ribosomal proteins and one endogenously biotinylated protein, as these are common contaminant proteins and are unlikely to be true KH interaction partners. More detailed experimental data are reported in Table S2 and S3 in Supporting Information.

Because proximity labeling typically identifies many proteins that are not bona fide partners of the BirA* fusion protein, we employed the complementary method of tandem affinity purification (TAP) to identify potential KH partners. KH is tightly bound to microtubules so that standard immunoprecipitation approaches would coprecipitate many microtubule-bound proteins that are not specifically associated with KH. To avoid this complication, we treated cells with formaldehyde to crosslink KH to partners that are within 2.3 – 2.7 Å (11) and then solubilized the parasites in strongly denaturing conditions so that only covalently crosslinked proteins would remain associated. In these experiments, KH was first fused to the HBH tag (11) that consists of a hexa-histidine motif, followed by a peptide that is a substrate for endogenous biotinylation, followed by another hexa-histidine motif. This TAP tag is designed to allow sequential purification under strongly denaturing conditions with a metal affinity resin followed by streptavidin resin. Using tandem mass tagging followed by tandem mass spectrometry, we identified proteins that were specifically enriched in the *∆kh*/HBH::KH formaldehyde-crosslinked sample as compared to a WT formaldehyde-crosslinked sample, thereby eliminating proteins that were abundant or interacting non-specifically with the metal affinity or biotin resins used for sequential affinity purification. Proteins that were enriched in the crosslinked sample in two duplicate experiments are listed in Fig. 1C, with likely contaminants, such as ribosomal proteins, excluded from the list. The proteins are listed in the order of their enrichment in Experiment 1, as determined by the difference in total protein intensity calculated from the TMT reporter ion intensities of peptide spectral matches for each protein. Their relative rank in the results is given for each experiment. Only three proteins were enriched in both repeats of the BioID and TAP experiments. One of them, α-tubulin, was expected, since earlier experiments had shown that KH was tightly associated with microtubules (4). The other two proteins are candidate KH partners and were named KH-Associated Proteins 1 and 2, or KHAP1 (LmxM.32.2440) and KHAP2 (LmxM.05.0380). The list of the top twenty identified proteins from each experiment can be found in tables S2 and S3.

### KHAP1 and KHAP2 predicted protein features

Bioinformatic analysis of KHAP1 indicates that this 502-amino acid, 56 kDa protein has orthologs broadly distributed among the kinetoplastid protists (tritrypdb.org/tritrypdb/). However, HMMER analysis (13) against the UniProtKB protein sequence database showed no significant homologies outside of kinetoplastid protists, suggesting that KHAP1is a kinetoplastid-specific protein. There were no conserved InterPro domains, but the NCBI Conserved Domain Database (14) detected a hit (E value of 9.37e-04) with the SMC-prok_B superfamily (cI37069) of SMC chromosome segregation proteins, although the significance of this low-scoring similarity is unclear. Interrogation of secondary structure using PSIPRED (15) predicted 59% helix content. Analysis using the COILS program (16) and a 21-amino acid window gave two strong predictions of coiled-coils between amino acids 72-103 and 114-134. The prediction of coiled-coils suggests that these regions of the protein could be involved in protein-protein interactions via such structures, although the sequence of KH is not predicted to have any coiled-coils using the COILS program.

KHAP2 is the ortholog of MARP-1 (microtubule-associated repetitive protein 1, Tb927.10.10360) from *T. brucei* that has been the subject of several previous studies in that parasite (17–19). This large 3196-amino acid, 374 kD protein is closely associated with the subpellicular microtubules, but not the axonemes, of trypanosomes, but specific biological functions for MARP-1 are not known. MARP-1 consists largely of 38-amino acid repeats (except for short unique N- and C-terminal domains) that are predicted to have both helical and non-helical segments. Due to the highly repetitive nature of the sequence of the orthologous *L. mexicana* KHAP2, the entire sequence of the ORF has not been determined from the *L. mexicana* genome, and the predicted ORF ends after only 783 amino acids, followed by undefined DNA sequence (tritrypdb.org/tritrypdb/), thus leaving the majority of the ORF unresolved. Nonetheless, the sequenced component of KHAP2 consists of similar 38-amino acid repeats and is presumably parallel in overall structure to MARP-1, consistent with our detection of several N-terminally epitope-tagged KHAP2 species of over 200 kDa mobility (see Fig 4C below). The reason for appearance of multiple species of distinct mobilities for both KHAP1 and KHAP2 in Western blots is not clear, but proteolytic degradation or post-translational modifications are possible explanations.

### KHAP1 and KHAP2 co-localize with KHARON at the subpellicular microtubules

To confirm potential association of each protein with KH, we epitope tagged them to test for overlap of fluorescent signal with KH. Immunofluorescence of Ty1::KHAP1 and OLLAS::KH (20) showed that KHAP1 colocalized with KH at the subpellicular microtubules (Fig. 2A). Analysis by Pearson’s Correlation Coefficient (CC) showed a positive correlation of 0.66. A single protein labeled with two different antibodies had a CC of 0.90 in the same analysis, and a negative control where the two channels are rotated 90 degrees to one another had a CC of. 02, as shown in supplemental figure S1. To eliminate the possibility of fixation artifacts, we endogenously tagged KHAP1 with the mNeonGreen fluorescent protein (mNG) (21). This fluorescent fusion protein also localized to the cell periphery in live cells (Fig. 2B,C). The localization was also performed for KHAP1 in amastigotes, detecting KHAP1::mNG by endogenous fluorescence in formaldehyde-fixed cells and colocalizing it with tubulin to show the amastigote periphery (Fig. 2D).

**Figure 2.**
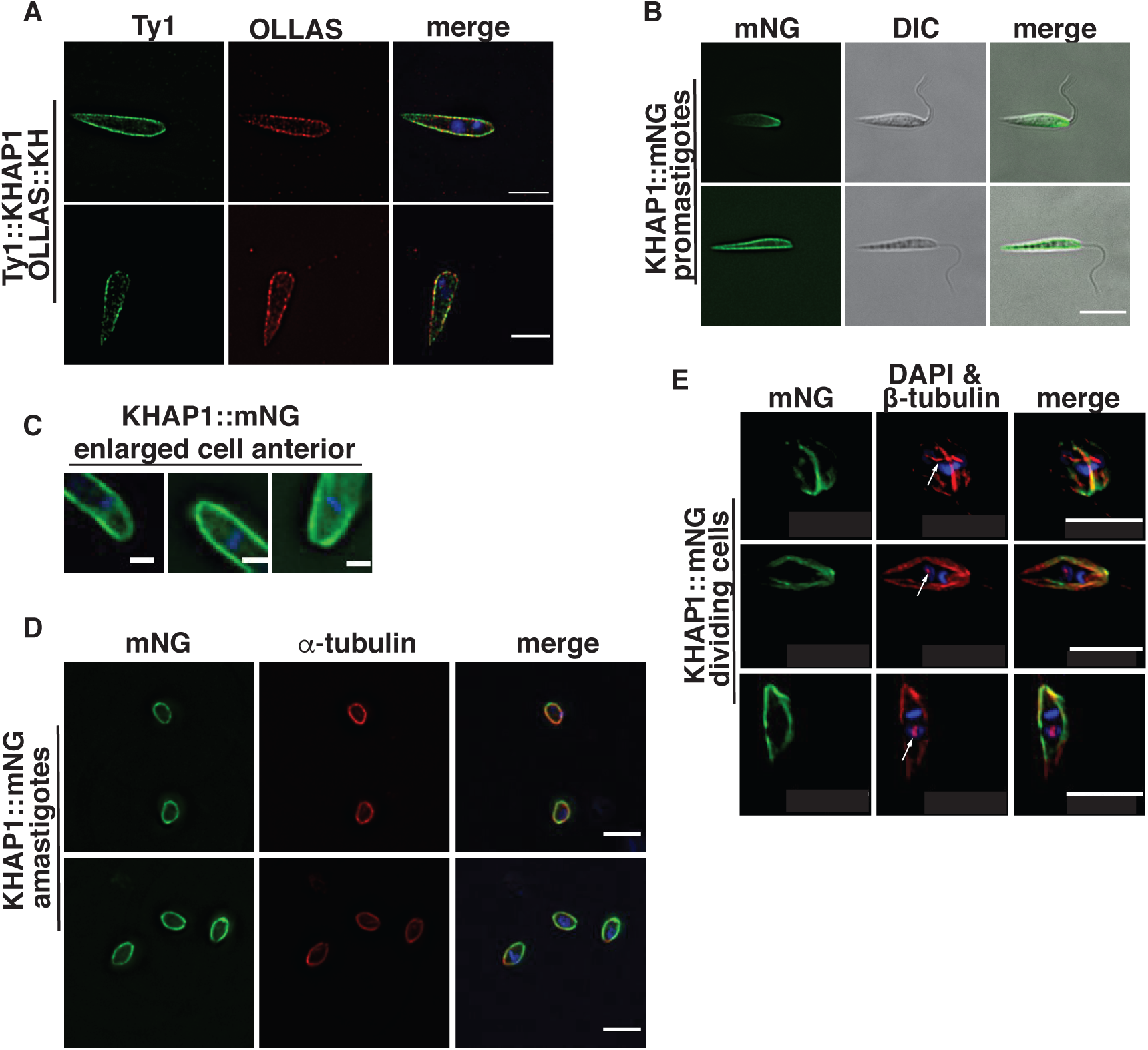
KHAP1 overlaps with KH at the sub-pellicular microtubules but not in the base of the flagellum or the mitotic spindle. *A*, Immunofluorescence of promastigotes expressing Ty1::KHAP1 and OLLAS::KH, probed with anti-Ty1 (*green*) and anti-OLLAS (*red*) antibodies, and stained with DAPI (*blue)*, scale bar 5µm. *B*, Live-cell imaging of promastigotes expressing fluorescent *KH*AP1::mNG suspended in CyGEL, scale bar 5 µm. *C*, Enlarged images of formaldehyde-fixed promastigotes showing mNG fluorescence (*green*) and DAPI staining (*blue*) focusing on the anterior regions of each parasite, scale bar 1 µm. *D*, Fixed amastigotes expressing KHAP1::mNG (*green*), stained with anti-α-tubulin antibodies (*red*) and DAPI (*blue*), scale bar 5 µm. *E*, Images of dividing promastigotes showing mNG fluorescence (green), β-tubulin stained with the *KMX-1* antibody (*red*), and the merged imaged (*merge*). The *white arrows* indicate the spindle in each cell that separates two lightly DAPI-staining nuclei. Strong blue stain represents kinetoplast DNA, scale bar 5 µm.

Immunofluorescence of Ty1::KHAP2 and KH, employing a rabbit anti-KH polyclonal antibody, showed that these proteins also colocalize at the cell periphery with a Pearson’s Correlation Coefficient of 0.58 (Figs. 3A, S1). In live cells mNG-KHAP2 also localized to the cell periphery (Figure 3B). In addition, KHAP2 localized to one to two small structures (yellow arrowheads, Figure 3A, 3B, and 3C) that are adjacent to the DAPI-stained kinetoplast DNA. Expanded images of mNG::KHAP2 expressing promastigotes (Fig. 3C) show these dot-like structures, close to the kinetoplast DNA, as well as a filament at the base of the flagellum. The dot structures are likely basal bodies, which are located adjacent to the kinetoplast DNA, and the filament is probably proximal regions of the axoneme. However, more detailed localization studies will be required to determine the precise distribution of KHAP2 in this region of the cell. Finally, mNG::KHAP2 also localized to the cell periphery in amastigotes, although in these smaller cells it is unclear whether mNG-KHAP2 still localized to the basal-body-like structures (Figure 3D). Notably, similar expanded images of cells expressing KHAP1::mNG (Fig. 2C) do not show fluorescence in the basal body or base of the flagellum.

**Figure 3.**
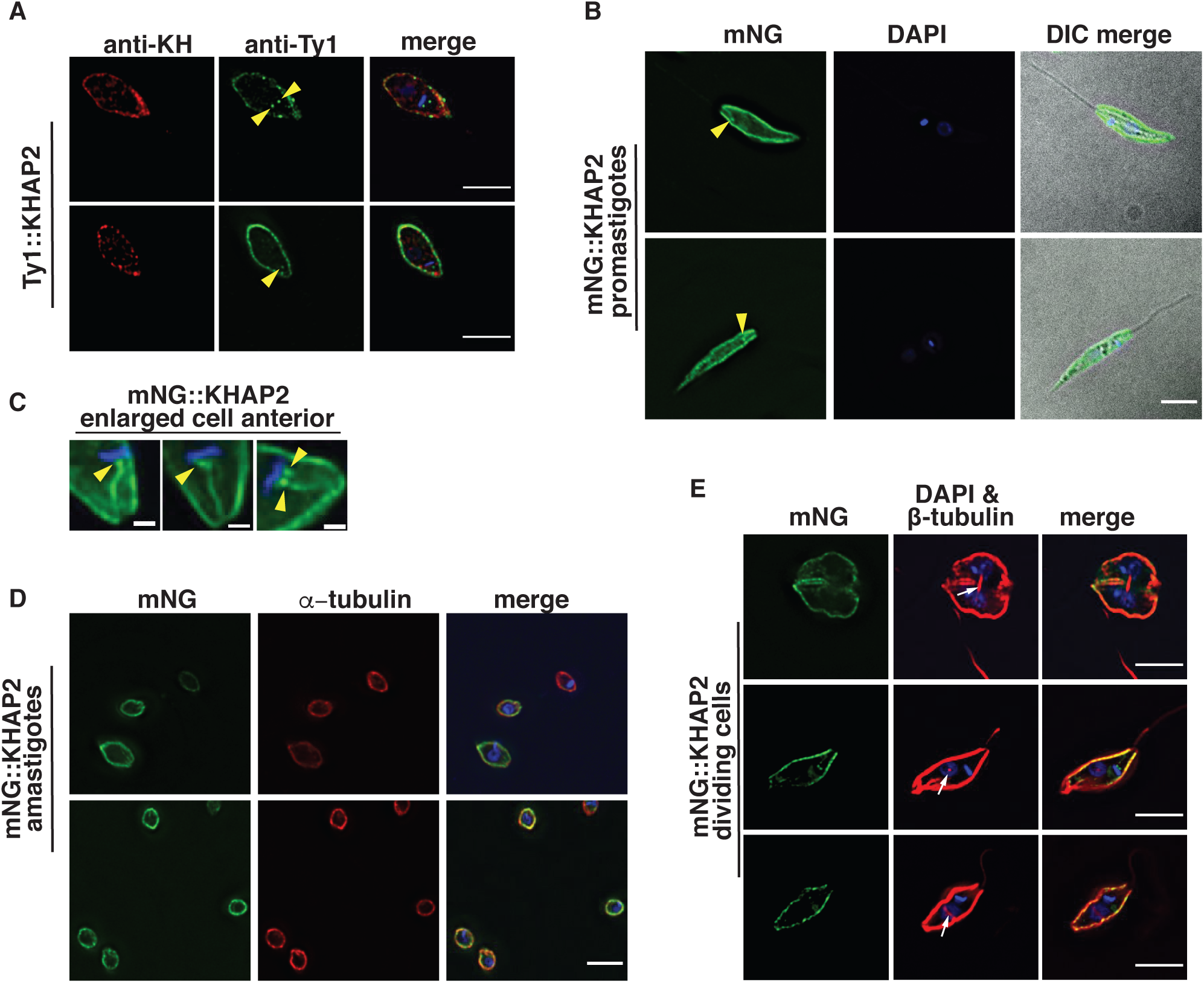
KHAP2 overlaps with KH at the sub-pellicular microtubules but not in the mitotic spindle. *A*, Immunofluorescence of promastigote expressing Ty1::KHAP2 probed with anti-Ty1 (*green*) and anti-KH (*red*) antibodies, and stained with DAPI (*blue*), *yellow arrowheads (A,B,C)* indicate spot-like structures adjacent to kinetoplast DNA here and in panels *B* and *C*, scale bar 5 µm. *B*, mNG::KHAP2 expressing promastigotes (*green*), fixed and stained with DAPI (*blue*), scale bar 5 µm. *C*, Enlarged images of formaldehyde-fixed promastigotes showing mNG fluorescence (*green*) and DAPI staining (*blue*) in the anterior region of each parasite, scale bar 1 µm. *D*, Fixed amastigotes expressing mNG::KHAP2 (*green*), stained with anti-α-tubulin antibodies (*red*) and DAPI (*blue*), scale bar 5 µm. *E*, Images of dividing promastigotes showing mNG fluorescence (green), β-tubulin stained with the *KMX-1* antibody (*red*), and the merged imaged (*merge*). The *white arrows* indicate the spindle in each cell that separates two lightly DAPI staining nuclei. Strong blue stain represents kinetoplast DNA, scale bar 5 µm.

### KHAP1 and KHAP2 do not localize to the mitotic spindle

Because one of the three locations observed for KH was the mitotic spindle (6), it was also important to determine whether KHAP1 or KHAP2 associated with KHARON at this site. Promastigotes expressing KHAP1::mNG were stained with the anti-β-tubulin KMX-1 antibody that allows visualization of mitotic spindles (22), and parasites in the process of division were imaged. Fig. 2E shows KHAP1::mNG (green) at the cell periphery but not at the centrally located β-tubulin (red, white arrows) that is positioned in between two DAPI-stained nuclei (light blue). In such images, the bright blue body is the duplicated kinetoplast DNA, whereas the nuclei stain less intensely with DAPI. Similarly, mNG::KHAP2 does not localize to the mitotic spindle (Fig. 3E).

Overall, KHAP1 overlaps with KH at the subpellicular microtubules but not at the base of the flagellum or in the mitotic spindle, while KHAP2 has a similar localization but is also likely present at the base of the flagellum. Hence, KHAP1 is selective for the subpellicular microtubules, or the postulated KH Complex 2 (Introduction), and can be employed to interrogate the function of that complex.

### KHAP1 and KHAP2 physically interact with KHARON as shown by formaldehyde crosslinking and pulldown

To verify that KHAP1 and KHAP2 are part of a complex with KH, we performed pulldown experiments under denaturing conditions, comparing formaldehyde-treated samples to untreated samples. In each case we used a His_10_-tagged KH (HIS::KH) as the bait for the pulldown, and tagged the protein of interest with the Ty1 epitope to enable its detection on Western blots. KH was detected with a rabbit antibody, generated in this study, which has strong specificity for KH (Fig. 4A). HIS::KH bound and eluted from the cobalt resin in both the crosslinked and uncrosslinked samples (Fig. 4B-D, upper panels marked HIS::KH, eluate, + crosslink and eluate, untreated), but both Ty1::KHAP1 and Ty1::KHAP2 were specifically enriched in the eluate only when the samples were crosslinked (Fig. 4B-C, lower panels marked Ty1::KHAP1 or Ty1::KHAP2). These results indicate that KHAP1 and KHAP2 are in close enough proximity to KH to be crosslinked by formaldehyde, but that the interaction is disrupted under denaturing conditions if no crosslinking is performed, (Fig. 4B and 4C). To demonstrate that the Ty1 tag did not independently associate with HIS::KH, we repeated the protocol with another protein, LmxM.29.2330, that was identified as a potential KH partner in the initial KH BioID experiment (Fig. 1B) but subsequently shown not to be a bona fide partner. Ty1-tagged LmxM.29.2330 was not enriched in the eluate in either the crosslinked or uncrosslinked condition (Fig. 4D), so the observed interactions between HIS::KH and Ty1::KHAP1 or Ty1::KHAP2 are not driven by the epitope tags on these proteins or other non-specific interactions.

**Figure 4.**
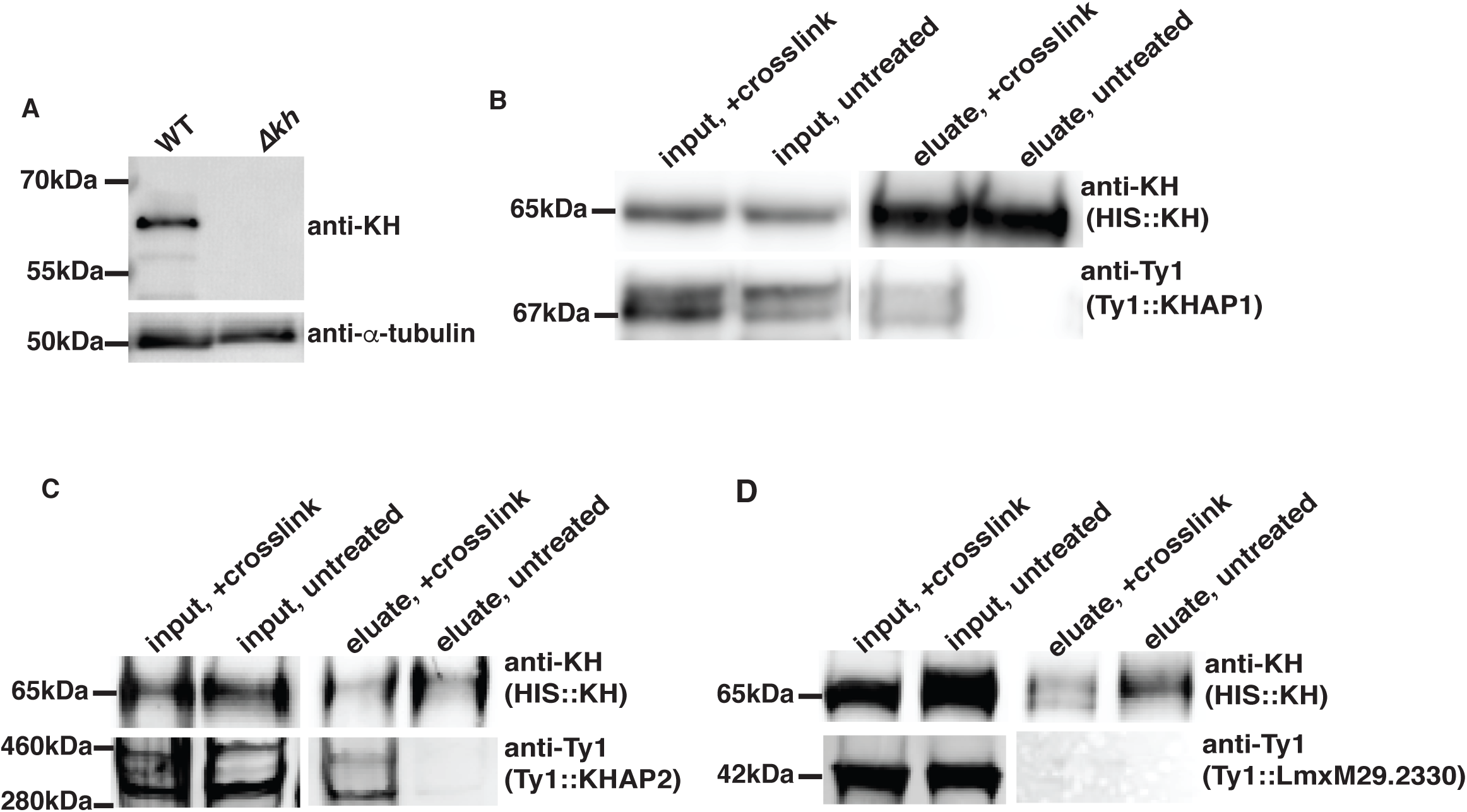
Association of KHAP1 and KHAP2 with KH determined by formaldehyde-crosslinking and pull downs. *A*, Characterization of the rabbit anti-KH antibody by immunoblot. Total lysates from both wild type (*WT*) and ∆*kh* null mutants were separated by SDS-PAGE, blotted onto a nitrocellulose membrane, and probed with the affinity purified anti-KH antibody or with anti-α-tubulin antibody, as indicated at the right. Numbers at the left indicate the positions of protein molecular weight markers in kDa. *B*, HIS::KH interaction with Ty1::KHAP1. Pulldown of HIS::KH under denaturing conditions with *(+ crosslink*) or without (*untreated*) formaldehyde crosslinking. Inputs and eluates from Co^2+^ affinity resin were separated by SDS-PAGE, blotted onto a nitrocellulose membrane, and probed with anti-KH and anti-Ty1 antibodies as indicated. In this and other panels, the text in parentheses below each antibody designation represents the protein being detected by the relevant antibody. *C*, HIS::KH interaction with Ty1::KHAP2. Pulldowns were performed as in panel *B*. *D*, Unrelated control protein, Ty1::LmxM29.2330, does not interact with HIS::KH. Pulldown as in panel *B*. In each blot, kDa numbers on the left indicate either the molecular weight markers that have a similar size to the protein of interest or the apparent size of the protein, calculated relative to the migration of two nearby molecular weight markers.

### Proximity ligation assays confirm close association of KH with KHAP1 and KHAP2

To further confirm the close association between KHAP1 or KHAP2 and KH, we performed the Proximity Ligation Assay (PLA) as an alternative test of molecular association (23,24). In this method, antibodies that recognize each of two interacting proteins are complexed with secondary antibodies that are covalently linked to oligonucleotides that can participate in rolling circle DNA amplification, provided that the two target proteins are located within ~ 40 nm of one another and can bring the two oligonucleotides close enough to allow their base pairing to additional connector oligonucleotides, followed by ligation of the connector oligonucleotides into a circular substrate. Following rolling circle DNA amplification, the amplicon is hybridized to a fluorescently labeled probe, resulting in fluorescent puncta within the cell if the two target proteins are in close proximity. To test for proximity between KH and KHAP1/2 we used Ty1-tagged KHAP proteins and the anti-KH antibody. As shown in Figure 5A, both Ty1::KHAP1 and Ty1::KHAP2 supported DNA amplification and resulted in fluorescent signal when anti-Ty1 and anti-KH antibodies were applied. As a standard negative control, we performed the PLA with only the anti-KH antibody and detected no amplification signal when no anti-Ty1 antibody was included (Figure 5A, bottom panel). To further confirm the specificity of the PLA, we chose another microtubule-bound protein to serve as a negative control. The *L. mexicana* ortholog*, Lmx*GB4 (LmxM.29.1950), of the *T. brucei* microtubule-binding protein GB4 (25), tagged with the Ty1 epitope, also localizes to the peripheral microtubules, with its localization biased towards the posterior end of the cell (Fig. 5B). However, antibodies against this fusion protein (Ty1::GB4) and KH did not result in a positive PLA signal (Fig. 5C), showing that even with both antibodies present, generation of a PLA signal requires close proximity of the two target proteins.

**Figure 5.**
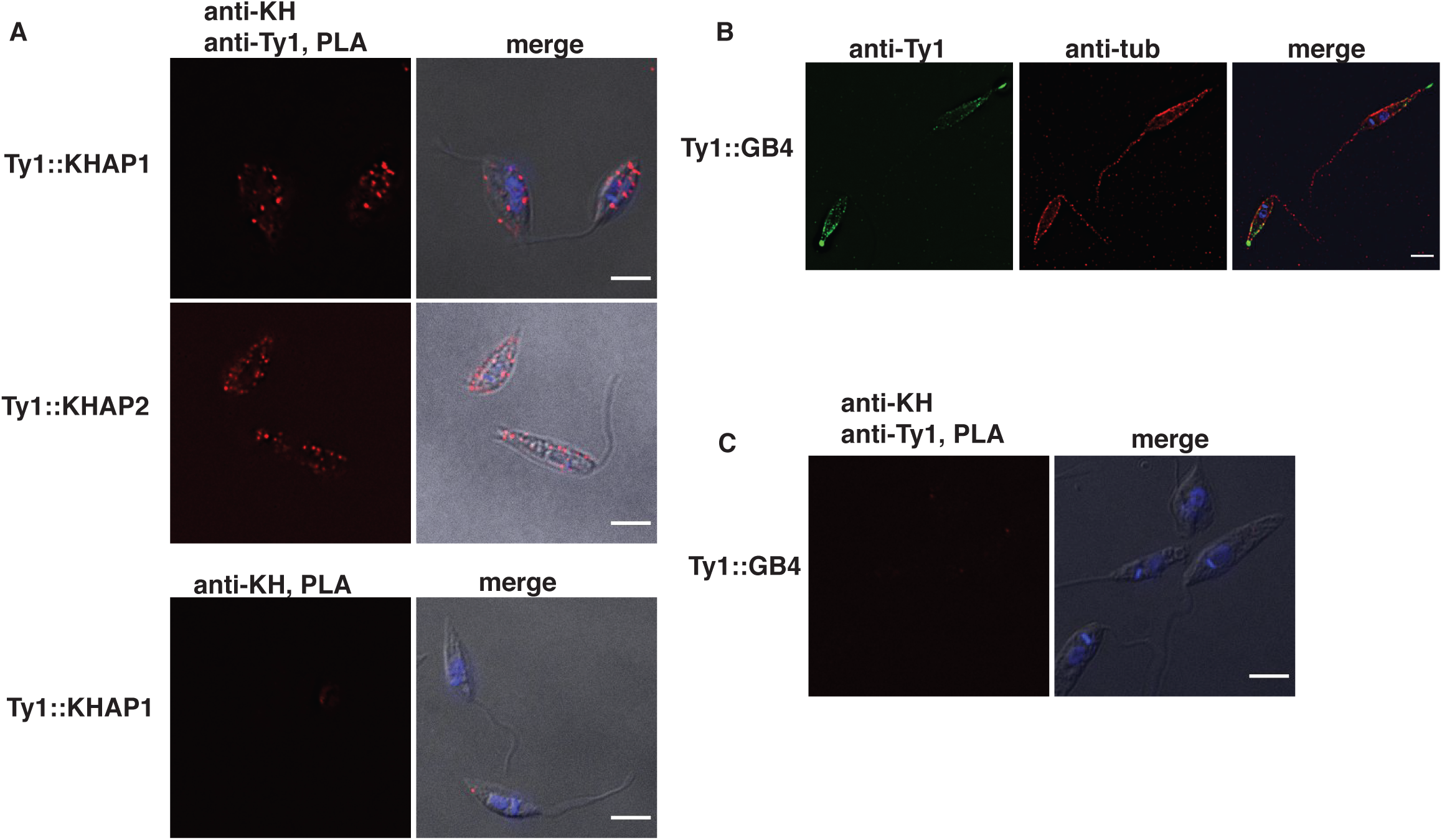
Proximity Ligation Assay (PLA) confirms the proximity of KHAP1 and KHAP2 with KH. Fluorescent signal indicating close proximity of KH with either KHAP1 (*Ty1::KHAP1*) or KHAP2 (*Ty1::KHAP2*) was detected only when both anti-KH and anti-Ty1 antibodies were employed (top and middle panel correspondingly). No fluorescent signal was detected when only anti-KH was used (bottom panel). Images marked *merge* show superpositions of the IF and DIC images. *B*, GB4 localizes to the cell periphery. IFA of Ty1::GB4 expressing parasites probed with anti-Ty1 and anti-α–tubulin antibodies. *C*, No PLA signal was detected between anti-KH and anti-Ty1 (Ty1-GB4-expressing parasites). Scale bars represent 5 µm.

### Generation of ∆khap1 and ∆khap2 null mutants

Since KHAP1 and KHAP2 are KH partner proteins that localized to the subpellicular microtubules, we next investigated whether they were involved in any of KH’s known cellular functions. To test KHAP functions, we generated null mutants of the *KHAP1* and *KHAP2* genes, using dual gene replacement with two drug selectable markers, mediated by CRISPR-Cas9 cleavage followed by homology-directed targeted gene replacement (26). The ∆*khap1* and ∆*khap2* null mutants were verified by PCR (Figs. 6A and B, respectively) to confirm that integration of each marker had occurred at the correct genomic location and that the relevant open reading frame (ORF) had been deleted. In addition, the ∆*khap1* null mutant was complemented with the *KHAP1* ORF on the pX63HYG (27) episomal expression vector to test for restoration of phenotypes. Since the complete *KHAP2* ORF has not been determined, the C-terminal coding sequence and 3’-UTR of this gene are not known. Therefore, we designed a knockout strategy employing an internal sequence within the repeat for targeting of the 3’ integration of each marker. Hence the PCR strategy for confirming the ∆*khap2* null mutant employed one primer that is upstream of the *KHAP2* gene and internal primers for each selectable marker. In addition, because the *KHAP2* ORF is not completely defined, we could not generate a similar episomal complement of the ∆*khap2* knockout. For this reason, we isolated a second independent ∆*khap2* clonal line and designated the two lines ∆*khap2.*c1 and ∆*khap2.*c2. These two lines allowed us to confirm phenotypes independently, thus supporting the conclusion that the phenotypes were due to deletion of the *KHAP2* gene. Typically, where the ∆*khap2* null mutants generated a phenotype different from wild-type parasites, we confirmed that difference with both clones, whereas phenotypes that were not changed by knockout of the *KHAP2* gene were investigated with the ∆*khap2*.c1 clone.

**Figure 6.**
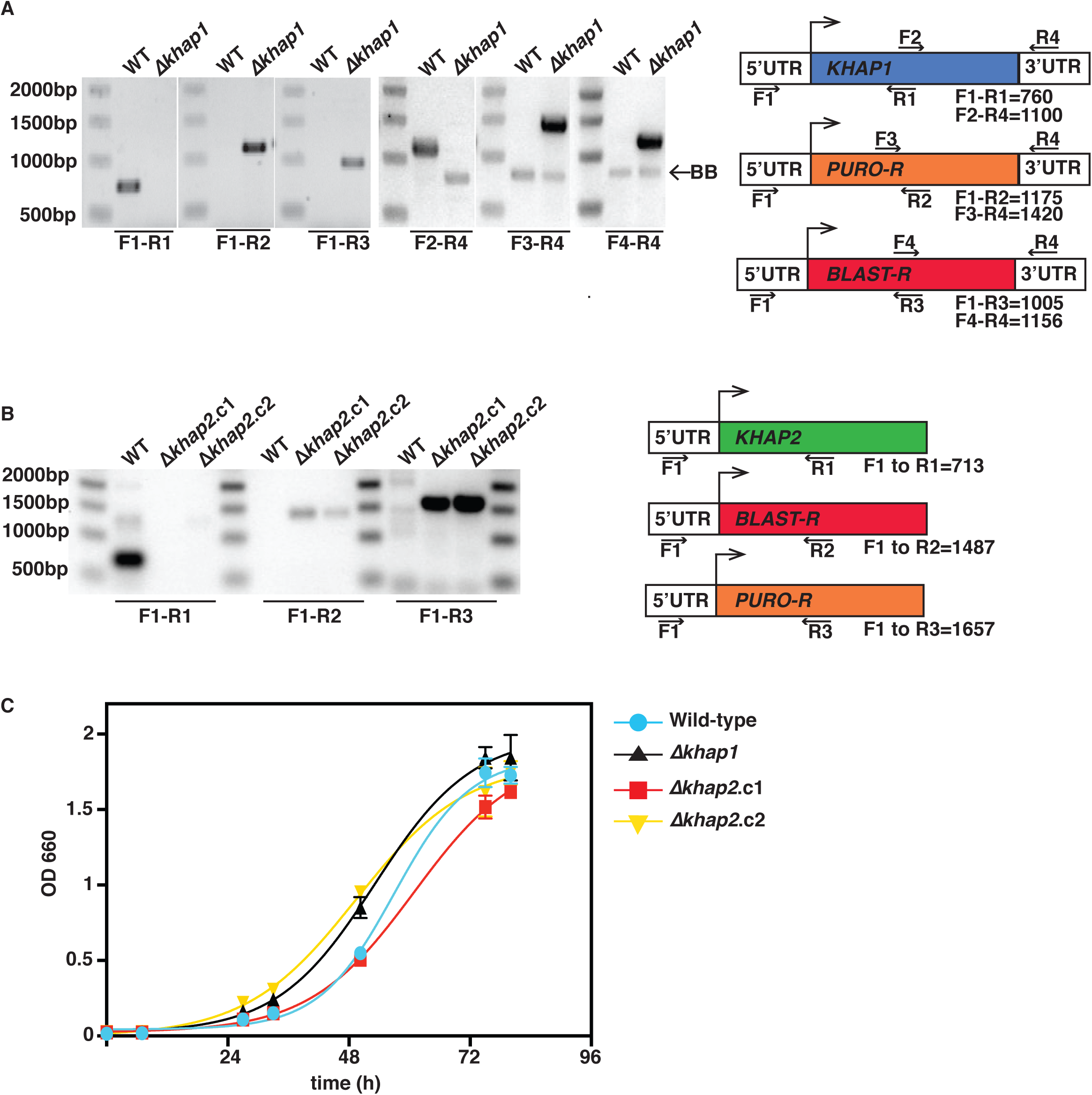
Confirmation of ∆*khap1* and ∆*khap2* null mutants and growth as promastigotes. *A*, PCR confirmation of the deletion of *KHAP1* by replacement of the ORF with the puromycin- and blasticidin-resistance genes. PCR amplification products demonstrate proper 3’ and 5’ integrations and loss of the *KHAP1* ORF. The primer pairs employed for each PCR are indicated below the relevant lanes. Not-to-scale diagram at the right indicates the structure of the gene locus, with colored boxes representing the relevant ORFs and white boxes representing the 5’- and 3’-untranslated regions (UTRs). Numbers underneath the gene locus indicate the predicted amplification products for the relevant primers in base pairs (*bp*). *BB* represents a background band amplified from both WT and ∆*khap1* DNA. *B*, PCR confirmation of the deletion of *KHAP2*, showing the replacement of the ORF by integration of puromycin- and blasticidin-resistance genes at the 5’ end of the gene. Due to the highly repetitive nature of the gene and surrounding sequence, the sequences of the C-terminal component of the ORF and the 3’ UTR are unknown. ∆*khap2*.c1 and ∆*khap2*.c2 represent two independent null mutants of this gene, as described in the text. *C*, Growth curve of WT, ∆*khap1* and the ∆*khap2*.c1 and *∆khap2*.c2 parasites. Growth was measured by OD_660_, and data points represent the average and standard deviations of three biological replicates per experiment.

### Phenotypic characterization of ∆khap1 and ∆khap2 null mutants in promastigotes

To evaluate the function of KHAP1 and KHAP2 and the KH Complex 2 containing these partners at the subpellicular cytoskeleton, we first examined phenotypes of the mutant in promastigotes. Neither mutant was compromised for growth in culture as promastigotes (Fig. 6C), an expected result given that the ∆*kh* null mutant is also not impaired in growth as promastigotes (4). KH was initially identified by its interaction with the glucose transporter GT1 and is required for efficient trafficking of GT1 to the parasite flagella. We tested whether KHAP1 or KHAP2 was required for KH’s role in the flagellar trafficking of GT1 by observing the localization of episomally expressed GT1::GFP in the Δ*khap1* and Δ*khap2* mutants. In each of these mutants the localization of GT1 was unaltered as compared to wild-type cells (Fig. 7A), as was confirmed by quantification of flagellar trafficking in each null mutant (Fig. 7B). These results indicate that these partner proteins are not required for the trafficking function of KH and imply that this KH complex located at the subpellicular cytoskeleton does not play a role in flagellar trafficking.

**Figure 7.**
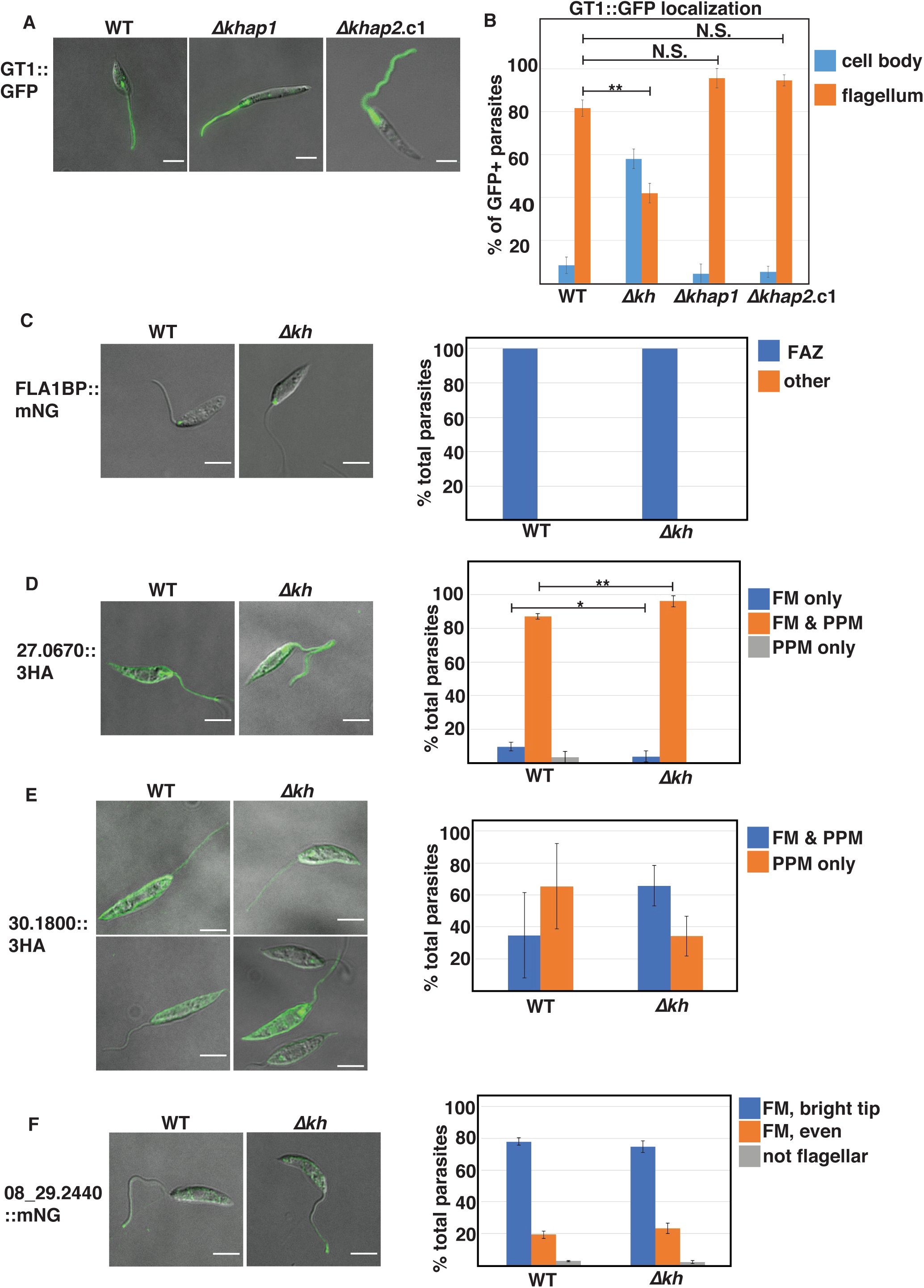
KHAP1, KHAP2, and KH are not involved in targeting of various proteins to the flagellar membrane. *A – B*, KHAP1 and KHAP2 are not required for GT1::GFP trafficking to the flagellar membrane. *A*, Live cell imaging of *WT* (*left panel*), *∆khap1* (*middle panel*), and ∆*khap2.*c1 (*right panel*) promastigotes episomally expressing GT1::GFP, imaged for GFP fluorescence and DIC in CyGEL. *B*, Quantification of GT1::GFP localization, with cells categorized by the localization of the bulk of fluorescent signal by an observer blind to the genetic strain. Data represent the average and 95% confidence intervals of three independent experiments. For each sample, between 25-40 cells were counted for each independent experiment. *C – F*, Four other membrane proteins do not depend on KH for flagellar targeting. *C*, FLA1 binding protein LmxM.10.0620, *FLA1BP*, *D*, putative amino acid permease LmxM.27.0670, *E*, putative amino acid permease LmxM.30.1800, *F*, putative cyclic nucleoside monophosphate phosphodiesterase LmxM.08_29.2440. For *C* and *F*, proteins were endogenously tagged at the C-terminus with mNG, whereas for *D* and *E*, proteins were tagged at the C-terminus with the 3HA tag and expressed from an episome. Panels at the *left* represent endogenous fluorescence (mNG) or immunofluorescence (3HA) for wild type (*WT*) parasites, and panels on the *right* are similar images for ∆*kh* null mutant lines. *Graphs* represent quantification of the *% total cells* with the indicated distribution of protein signals, for *WT* or ∆*kh* parasites, as determined by microscopic examination of between 48 and 428 parasites per sample in three replicate experiments. Error bars represent 95% confidence intervals. Statistical significance was determined between pairs using Student’s t-test, with *N.S.* indicating not significant, * representing p<0.05, and ** p<0.01. For *C-F* all comparisons not marked by asterisks were not significantly different. Scale bars represent 5 µm. Abbreviations used in panels at the right are: FAZ, flagellum attachment zone; FM, flagellar membrane; PPM, pellicular plasma membrane.

### KH is not required for trafficking of several other membrane proteins to the flagellum

While KH was initially identified as a protein important for efficient trafficking of GT1 to the flagellar membrane, it is unclear how extensive a role this protein plays in assembly of the flagellar membrane. To address this question, we studied in wild type and ∆*kh* null mutants the trafficking of four other proteins that are present in the flagellar membrane. The FLA1 binding protein, FLA1BP or LmxM.10.0620, is a single membrane pass protein that is in the flagellar membrane component of the flagellum attachment zone (FAZ) (28), a discrete adhesion between the flagellar and flagellar pocket membranes in the proximal region of the flagellum (Fig. 7C). A putative cyclic nucleotide phosphodiesterase, LmxM.08_29.2440, is a single membrane pass protein detected in the flagellum by proteomics (29) and shown here to be in the flagellar membrane with an accumulation at the distal tip (Fig. 7F). Two putative amino acid transporters also detected in the flagellar membrane in the proteomic study, LmxM.27.0670 and LmxM.30.1800, distribute between the pellicular plasma membrane and the flagellar membrane (Fig. 7D,E). Fluorescence microscopy and quantitative analysis of such images (Fig. 7C-F) establish that each of these proteins traffics equally well to the flagellar membrane in wild type and ∆*kh* null mutant promastigotes and that KH is thus not required for efficient flagellar trafficking of several integral membrane proteins with diverse localization patterns in this organellar membrane. There are some small but statistically significant differences in Fig. 7D, but the data do not invalidate the conclusion that KH plays little if any role for FM trafficking of these proteins.

### KH association with the cytoskeleton is independent of KHAP1 and KHAP2

We also interrogated the relationship of KH, KHAP1, and KHAP2 to one another in terms of potential dependency for subcellular localization. Using Ty1::KHAP1, we first tested whether KHAP1 required KH for its localization to the subpellicular microtubules. The localization of Ty1::KHAP1 was unchanged in the absence of KH (Fig. 8A). Similarly, deletion of the *KHAP1* ORF does not alter the localization of KH to the cell periphery (Fig. 8B). Furthermore, deletion of the *KHAP2* ORF did not alter the localization of HA::KHAP1 to the cell periphery (Fig. 8C). Finally, fractionation of the ∆*khap1* and ∆*khap2* null mutants into detergent soluble (S, supernatant) and insoluble (P, pellet) components showed that neither KH partner is required for KH association with the cytoskeletal pellet (Fig. 8D). Hence, these partner proteins are not co-dependent upon each other for cytoskeletal association.

**Figure 8.**
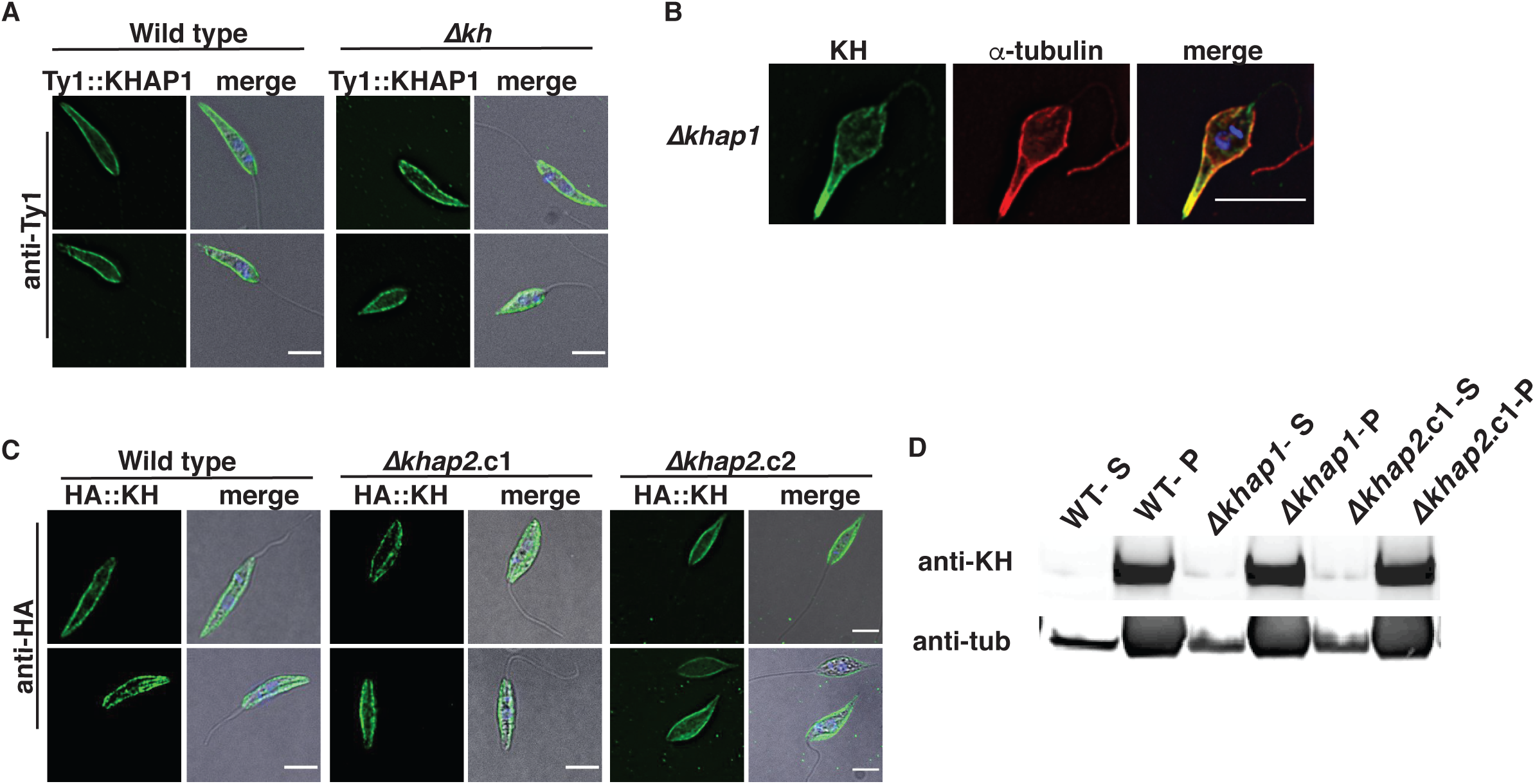
KH, KHAP1, and KHAP2 show independent cytoskeletal association. *A*, Immunofluorescence of wild-type and *∆kh* cells expressing Ty1::KHAP1, probed with anti-Ty1 (*green*) and stained with DAPI (*blue*), merged with DIC in the right panel. *B*, Immunofluorescence of *∆khap1* cells probed with anti-KH (*green*) and anti-α-tubulin (*red*) and a merged image with DAPI staining (*blue*) and DIC at the right. *C*, Immunofluorescence of wild-type and *∆khap2* cells expressing HA-KH, probed with anti-HA (*green*), merged image with DAPI staining (*blue*) and DIC in the right panel. *D*, Western blot of cytoskeleton-bound proteins from parasites of indicated genotypes, isolated by extraction with 1% NP-40 and centrifugation. Detergent-soluble and cytoplasmic proteins are found in the supernatant (*S*), and cytoskeleton-bound proteins are found in the pellet (*P*). Blot probed with anti-KH and anti-α-tubulin (*anti-tub*) antibodies. Scale bars represent 5 µm.

### Phenotypic characterization of ∆khap1 and ∆khap2 null mutant in amastigotes

One striking phenotype of ∆*kh* null mutants is their inability to survive as amastigotes following entry into mammalian macrophages. These mutant amastigotes fail to execute cytokinesis, generating multi-nucleated, multi-flagellated parasites (4), and this deficiency also makes the ∆*kh* null mutants avirulent in mice (5). To examine the roles in amastigotes of KHAP1, KHAP2, and hence the KH Complex 2 at the subpellicular cytoskeleton, we infected THP-1 macrophages with the ∆*khap1* null mutant and its complemented line and with the two independently generated ∆*khap2* null mutant lines and quantified the number of remaining intracellular amastigotes at days 1, 4, and 7 after infection (Fig. 9A,D, respectively). The ∆*kh*, ∆*khap1*, and ∆*khap2* null mutants infected macrophages at day 1 as well as wild type parasites, but while wild type amastigotes replicated robustly by day 7, the null mutants did not. Complementation of the ∆*khap1* null mutant by episomal expression of the *KHAP1* ORF partially restored the wild type phenotype, confirming that the impairment of amastigote replication was due to deletion of this gene. For the ∆*khap1* null mutants, we also investigated the occurrence of multinucleated amastigotes. Similar to Δ*kh* mutants, the majority of Δ*khap1* mutants had greater than 1 nucleus (1N), and >40% had more than two nuclei, with a striking increase in parasites with more than two nuclei and expanded DNA masses (Fig. 9B and C). Given that wild-type parasites were >80% 1N, this is evidence of a pronounced cell division defect. Microscopic examination revealed cells with several nuclei, and the DNA masses appear misshapen (Fig. 9C). The defect in amastigote cell division was genetically complemented when KHAP1 was episomally expressed as an add-back, indicating that the loss of KHAP1 was solely responsible for the defect. This increase in cells with >2 nuclei was also observed in *∆khap2* amastigotes (Fig. 9 E and F). Significantly, these results imply that the subpellicular cytoskeletal KH complex that contains KHAP1 and KHAP2 is required for cytokinesis and replication of intracellular amastigotes and is hence responsible, at least in part, for the avirulent phenotype previously defined for ∆*kh* null mutants.

**Figure 9.**
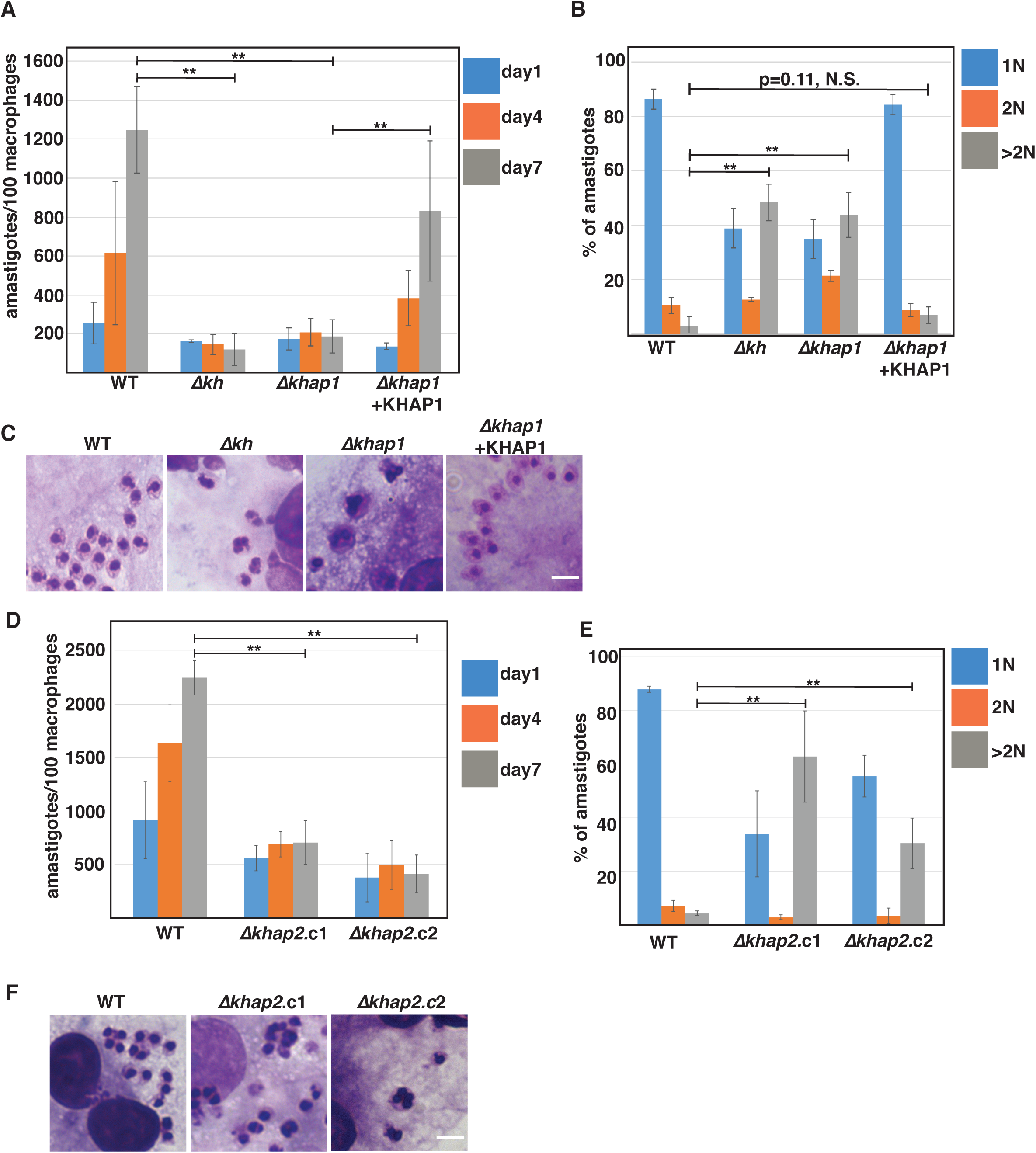
KHAP1 and KHAP2 are required for amastigote growth. *A*, KHAP1 is required for amastigote replication in THP-1 macrophages over seven days at 34.5°C. Parasite genotypes are indicated below the x-axis. Data represent the averages of three independent experiments with 100 macrophages assessed in each experiment. All error bars in this figure represent 95% confidence limits. Horizontal bars indicate data for which statistical significance was calculated by Prism 7 (GraphPad) using Student’s t-tests and a Holm-Šídák test to compute multiplicity adjusted p values for multiple comparisons, with ** representing p<0.01. *N.S.* indicates not significant. *B*, Number of nuclei per amastigote at day 4 of infection. The number of nuclei was estimated by microscopic examination of Giemsa-stained infected macrophages, with >100 amastigotes assessed in each experiment, three independent experiments per sample. Parasite genotypes are indicated below the x-axis. *C*, Giemsa-stained amastigotes growing within THP-1 macrophages show multiple nuclei in ∆*kh* and ∆*khap1* amastigotes. *D*, KHAP2 is required for amastigote replication in THP-1 macrophages. Experiments were performed as in part *A* employing ∆*khap2.*c1 and ∆*khap2.*c2. *E*, Number of nuclei estimated by microscopic examination for *WT*, ∆*khap2.*c1 and ∆*khap2.*c2 amastigotes. Experiments were performed as in part *B*. *F*, Amastigotes of ∆*khap2.*c1 and ∆*khap2*.c2 exhibit multiple nuclei. Scale bars represent 5 µm.

## Discussion

The cytoskeleton of *Leishmania* and trypanosomes is complex and consists of multiple microtubule-based structures, including the flagellar axoneme, the subpellicular microtubule cage or pellicular cytoskeleton that underlies the plasma membrane, and the mitotic spindle. Previous studies on the *L. mexicana* flagellar glucose transporter GT1 established that a cytoskeletal protein, designated KH, interacts with the flagellar targeting motif of that permease and is involved in trafficking GT1 to the flagellar membrane (4). KH in both *L. mexicana* and *T. brucei* is distributed over microtubule structures at the base of the flagellum, the pellicular cytoskeleton, and the mitotic spindle (4,6). Furthermore, studies with either gene knockouts in *L. mexicana* (4,5) or RNAi in *T. brucei* (6) established that KH is required for trafficking of some proteins to the flagellar membrane, for viability of infectious stage parasites, and for cytokinesis. Thus, KH is a multifunctional protein with multiple subcellular locations. Deconvolving multiple functions for a protein with diverse subcellular locations can be challenging, as mutations in the cognate gene, especially gene knockouts, often affect all functions. One postulate of the current investigation was that KH is associated with multiple partner proteins, some of which may be unique to specific subcellular sites. Hence, we have advanced the hypothesis that KH may be associated with three discrete types of complexes located at distinct sites and with diverse functions: KH Complexes 1 (base of the flagellum), 2 (subpellicular microtubules), and 3 (mitotic spindle). According to this hypothesis, identifying partners unique to each complex will help resolve the diverse functions of KH, as deletion of genes encoding unique partners may compromise one complex but not the others. In the current investigation, we have identified two *L. mexicana* proteins, KHAP1 and KHAP2, that are members of KH Complex 2 at the subpellicular microtubules. Null mutants in each of these genes impair replication of intracellular amastigotes, blocking cytokinesis in the infectious stage of the parasite life cycle, but these mutants are not impaired in flagellar trafficking of GT1 in insect stage promastigotes. These results confirm that partner proteins specific to one subcellular site or KH Complex do exist, especially KHAP1 that is only found in the subpellicular microtubules, and that deletion of the cognate genes affects only a subset of the known KH functions, in this case the ability to execute cytokinesis in the amastigote stage. KHAP2 also appears at the base of the flagellum (Fig. 3C), although the ability of the ∆*khap2* null mutant to correctly target GT1 to the flagellar membrane (Fig. 7A, B) indicates that KHAP2 at this site is not required for targeting membrane proteins to the flagellum.

KHAP1 and KHAP2 were initially identified as high probability candidates for KH partners on the basis of both BioID and TAP. Their association with KH has been confirmed by i) pulldown of each KHAP with HIS::KH following formaldehyde crosslinking and metal affinity chromatography under strongly denaturing conditions, ii) positive signals between KH and each KHAP in the PLA, iii) overlap of fluorescence signals in light microscopy. Among these approaches, the most stringent is pulldown following formaldehyde crosslinking, as covalent crosslinks with formaldehyde have been estimated to require side chains of the relevant proteins to approach within ~2 Å of each other (30). The results from IFA and PLA provide orthogonal strategies that further support the association of KHAP1 and KHAP2 in complexes with KH.

KHAP1 appears to be a kinetoplastid-specific protein whose major predicted feature is two coiled-coil domains, but there are no predicted biological functions that can be discerned from the sequence. The coiled-coil domains could mediate interactions with other proteins or an internal association of the two structures. KHAP2 is the ortholog of the large microtubule-associated repetitive protein MARP-1 from *T. brucei*. While the association of the trypanosome protein with the subpellicular microtubules has been documented in previous studies, the specific function within the pellicular cytoskeleton is not well defined. The present work implicates KH, KHAP1, and KHAP2 as required for cytokinesis of amastigotes. It seems likely that KHAP1, KHAP2, and KH are components of the same Complex 2, but their precise arrangement and the potential existence of other currently unknown components is uncertain.

The observation that knockouts of the *KH*, *KHAP1*, and *KHAP2* genes all impair cytokinesis in amastigotes but not promastigotes implies that the pellicular cytoskeleton plays distinct roles in cell division in these two life cycle stages. We do not currently know how these null mutants block cytokinesis at a mechanistic level. However, a body of work in *T. brucei* establishes that the pellicular cytoskeletal network is extensively remodeled during cell division, including establishment of the cleavage furrow between the old and new flagella of the replicating parasite (e.g., (31–33)). One possibility is that the KH Complex 2 is required, directly or indirectly, for initiation or progression of the cleavage furrow in amastigotes, thus allowing intracellular parasites of the null mutants to undergo replication of nuclei and flagella but not to separate into daughter cells. As predicted, this inability to undergo cell division results in amastigote death *in vitro* and avirulence in a murine model of cutaneous leishmaniasis (5), hence underscoring the importance of KH and its associated Complex 2 proteins in the biology and virulence of these parasites.

Since the majority of KH is associated with the pellicular cytoskeleton, it is not surprising that the KH partners most readily identified are components of Complex 2. Efforts are now ongoing to define potential partners of the postulated KH Complexes 1 and 3. Recent studies employing BioID in *T. brucei* have identified KH as a proximity partner for each of five spindle-associated proteins (SAPs) designated NuSAP1, NuSAP2, Kif13-1, *Tb*Mlp2, and *Tb*AUK1 (34). These proteins play various roles in mitosis, including faithful chromosome segregation, G2/M transition, stabilization if kinetochore proteins, etc. Hence, orthologs of these proteins in *L. mexicana* are candidates for KHAPs at the mitotic spindle and could be investigated by approaches similar to those used here.

It is notable that two of the proteins identified by BioID as proximity partners for KH (Fig. 1B) are likely to be localized to the basal body. LmxM10.1150 is a homolog of the cilia- and flagella-associated protein 45 (35) and is the ortholog of Tb927.8.4580, a protein that has been localized to both the basal body (36) and the flagellum (tryptag.org) of procyclic trypanosomes. LmxM31.0380 is the ortholog of Tb927.10.14520, a basal body protein designated TbBBP52 (36). Hence, both hits are candidates for components of KH Complex 1 that are worthy of further detailed investigation. A complementary approach to identifying Complex 1 components would be to search for KH partners in isolated flagella, e.g., by TAP. Isolating flagella from *Leishmania* promastigotes is problematic, because such methods typically leave the base of the flagellum associated with the cell body (29) and thus would not be expected to separate Complex 1 components from those of the other KH Complexes. However, flagella or their component cytoskeletons can be isolated from *T. brucei* with the kinetoplast DNA, basal body, and proximal regions intact (37-39). Hence, that parasite offers a potentially tractable model for identifying KH Complex 1 components, and orthologs in both *T. brucei* and *L. mexicana* could be tested for roles in transport of flagellar membrane proteins such as *Tb*CaCh/FS179, a putative Ca^2+^ channel in *T. brucei* that is dependent upon *Tb*KHARON for flagellar localization (6), and GT1 (4), respectively.

The observation that four other integral membrane proteins are not dependent upon KH for flagellar trafficking (Fig. 7C – F) implies that other flagellar membrane trafficking machines likely exist that are independent of KH and its associated proteins. Given the multiplicity of vital biological roles played by flagellar membrane proteins in kinetoplastid parasites (40), these putative alternate trafficking machineries are worthy of investigation in their own right.

## Experimental Procedures

### Parasite culture and transfections

Wild type *Leishmania mexicana* MNYC/BZ/62/M379 promastigotes were cultured in RPMI 1640 medium (Gibco) supplemented with 10% heat-inactivated fetal bovine serum (FBS) (Thermo Scientific Hyclone, Logan, UT), 0.1 mM xanthine, 5 µg/ml of hemin, 100 units/mL penicillin (Gibco), and 100µg/ml streptomycin (Gibco). Promastigote cultures were maintained at 26°C without supplemental CO_2_. Relevant drugs for the selection of genetically modified parasites were added at the following concentrations: hygromycin B at 100 µg/ml, neomycin (G418) at 100 µg/ml, phleomycin at 20 µg/ml (InvivoGen), blasticidin at 25 µg/ml (InvivoGen), and puromycin at 10 µg/ml (InvivoGen).

For the introduction of the episome to express Ty::KHAP1 and the endogenous tagging of OLLAS-KH and HBH-KH, *L. mexicana* promastigotes were transfected according to previously described electroporation techniques using a Bio-Rad Gene Pulser Xcell (41,42). For the introduction of DNA for CRISPR/Cas9 experiments, including the episome encoding the Cas9 and T7 RNA polymerase enzymes, as well as the episomes for the expression of HIS::KH, GT1::GFP, Ty1::LmxM29.2330, 3xHA::LmxM.27.0670 and 3xHA::LmxM.30.1800, cells were transfected using an Amaxa Nucleofector II (Lonza) by administering a single pulse with program X-001 as previously described (43). To compare promastigote growth of wild-type, *∆khap1*, and *∆khap2* parasites, cultures were inoculated at 5×10^5^ cells/ml and cell density was monitory by optical density at 660 nm using a Pharmacia Biotech Ultraspec 2000.

### Genetic manipulations of parasites

Deletion mutants of the *KHAP1* and *KHAP2* genes were generated by CRISPR/Cas9 mediated gene replacement, as previously described (26), using the primer generation software at LeishGEdit (http://www.leishgedit.net). For each deletion mutant we used two drug cassettes to replace both alleles of the gene, encoding puromycin and blasticidin resistance genes. The published genomic sequence for the *L. mexicana* KHAP2 gene is incomplete due to the extremely repetitive nature of the ORF, so a primer was generated that targeted the repeated sequence in order to generate a interrupted open reading frame when paired with the LeishGEdit-generated 5’ targeting primer (*KHAP2*-repeat targeting sequence: GGCGGGGTCCACGGGCACCTTCTCGTAGCC). The following tagged lines were also generated by CRISPR/Cas9, as described for the deletions above, and the protein tags were selected for using puromycin-resistance: KHAP1::mNG, mNG::KHAP2, Ty1::KHAP2, Ty1::GB4.

To generate the endogenously-tagged OLLAS::KH and HBH::KH parasite lines we used the gene replacement technique described in (44). To construct OLLAS::KH, the donor plasmid encoded a cassette consisting of a blasticidin resistance marker, a self-cleaving viral TaV2A peptide (45), and the 3xOLLAS peptide tag (20). Flanking 5’ and 3’ targeting sequences were generated by PCR, then the donor vector and the targeting sequences were ligated together to form a targeting plasmid, digested with Swa1, the resulting targeting construct was transfected into wild-type parasites. Stably transfected parasites were selected by drug resistance. To construct HBH::KH, a donor plasmid was constructed encoding a cassette consisting of a blasticidin resistance marker, a self-cleaving viral TaV2A peptide (45), and the His_6_-biotinylation motif-His_6_ tandem-affinity purification tag (4,11), and then that construct was introduced using the same protocol as for OLLAS::KH. The HBH::KH construct was transfected into heterozygous wild-type/*∆kh* parasites so that the HBH::KH is the sole copy of KH present.

### Construction of episomes for protein expression

BirA*::KH was constructed by the PCR amplification of the Myc-BirA* cassette from the pLew100-Myc-BirA* expression vector (46), followed by subcloning into the Sma1 site of the *Leishmania* pX72-Hyg expression vector, which is a modified version of the pX63-Hyg expression vector (27)). Then the genomic KH open reading frame was PCR amplified and cloned into the Kpn1 and EcoRV sites in pX72-Hyg to generate pX72-Hyg-Myc-BirA*::KH. This episome was transfected into wild-type and *∆kh* parasites and BirA*::KH expression was verified by Western blot.

To generate Ty1::KHAP1 and Ty1::LmxM29.2330 we first subcloned the Ty1-epitope (47) into the pX63-Neo expression vector. Then KHAP1 and LmxM29.2330 ORFs were amplified from genomic DNA and subcloned into the Mfe1/Xba1 sites of the pX63-Neo-Ty1. Similarly, to create HIS::KH we first subcloned the 10xHIS tag, generated by annealing two long overlapping oligonucleotides, into the pX63-Neo expression vector using the EcoR1 site in the polylinker, then the KH ORF was PCR amplified from genomic DNA and cloned it into the Mfe1 and Xba1 sites of pX63-Neo-HIS to generate pX63-Neo-HIS::KH.

To generate LmxM.27.0670:: 3xHA and LmxM.30.1800:: 3xHA the ORFs for each gene were amplified from genomic DNA and subcloned into the Xma1 site of the plasmid pX63-Hyg-HA_3_, which was derived from pX63-Neo-3HA (9).

### BioID MS experiment

*L. mexicana* promastigote cultures of wild type parasites and the BirA*::KH-expressing parasites were grown to ~5 × 10^6^ cells/ml, then biotin was added to the cultures to a final concentration of 50 µM and cells were incubated overnight to a final density of 1-2×10^7^ cells/ml. Cells (1-2×10^9^) incubated with biotin were collected, washed twice with phosphate-buffered saline (PBS: 137 mM NaCl, 2.7 mM KCl, 10 mM Na_2_HPO_4_, 2 mM KH_2_PO_4_, pH 7.4), and then resuspended in PEME + protease inhibitors (2 mM EGTA, 1 mM MgSO4, 0.1 mM EDTA, 0.1 M piperazine-*N*,*N*′-bis(2-ethanesulfonic acid)–NaOH (PIPES-NaOH), pH 6.9 supplemented with cOmplete EDTA-free protease inhibitor cocktail (Roche Diagnostics)), and pelleted again. Detergent-extracted cytoskeletons were prepared by incubating the biotin treated cells in PEME buffer with protease inhibitors plus 0.5% Nonidet P-40 (Sigma) for 30 min at room temperature followed by centrifugation at 3400 × g for 5 min at room temperature. This pellet was then solubilized in lysis buffer (500 mM NaCl, 5 mM EDTA, 1 mM DTT, 50 mM Tris, 0.4% SDS, pH 7.4) by incubation on a rocker for 15 min followed by sonication using 3 pulses of 15 seconds at maximum power (Sonic Dismembrator, 500W, Fisher Scientific). After sonication samples were spun at 16,000 × g for 20 min at 4°C and the supernatant was added to 400 µl of streptavidin resin (Pierce® High Capacity Streptavidin-agarose Resin, Thermo Scientific Pierce) and incubated overnight at 4°C.

These streptavidin resins were then repeatedly washed by rotating mixing, 5 min for each wash. This included five washes with 1 ml of Buffer 3 (8 M urea, 200 mM NaCl, 100 mM Tris, pH 8.0) containing 0.2% SDS (5 min each), five washes with 1 ml of Buffer 3 with 2.0% SDS, five washes with 1 ml of Buffer 3 with no SDS (5 min each), two washes with 1 ml of Buffer A (200 mM NaCl, 100 mM Tris, pH 7.0), and finally two washes with 1 ml of Tris, pH 8.0. Samples were stored in 500 µl of Tris, pH 8.0, at 4 °C until trypsin digestion.

Trypsin digestion and analysis of tryptic peptides by mass spectrometry is detailed in Supporting Information and a summary of the data is presented in Table S1.

### Tandem affinity purification of HBH::KH

*L. mexicana* wild-type and *∆kh*/HBH::KH promastigotes were grown to a density of −2 × 10^7^ cells/ml. Approximately 1.5 × 10^9^ cells were collected, washed once with PBS, and resuspended in PBS. Cells were then cross-linked with 1% formaldehyde (Ultra Pure EM grade, Polysciences, Warrington, PA), for 10 min at 26°C followed by another 5 min incubation after adding glycine to a final concentration of 125 mM to stop the cross-linking reaction. PBS was added in place of formaldehyde for non-cross-linked control samples. Cells were washed once with PBS, then resuspended in 1 ml of Buffer 1 (8 M urea, 300 mM NaCl, 0.5% Nonidet P-40, 50 mM NaH_2_PO_4_, 50 mM Tris, pH 7.0) on ice. Samples were sonicated 3 times on ice at 50% max amplitude for 10 sec with 30 sec between pulses. Two percent of this protein lysate was saved as the protein input.

The remainder of the protein was incubated with 750 µl of HisPur^TM^ Cobalt Resin (Thermo Scientific Pierce, Rockford, IL) on a rocker for 45 min at room temperature. The resins were washed five times with 1 ml of Buffer 1 (5 min each), five times with 1 ml of Buffer 1 at pH 6.4 (5 min each), and five times with 1 ml of Buffer 1 pH 6.4 with 10 mM imidazole (5 min each). Cobalt-bound protein complexes were eluted with 1 ml of Buffer 2 (Elution buffer: 45 mM NaH_2_PO_4_, 8 M urea, 270 mM NaCl, 150 mM imidazole). The eluates were incubated with 400 µl of Pierce® High Capacity Streptavidin-agarose Resin (Thermo Scientific Pierce) on a rocker overnight at 4 °C. Streptavidin resins were then washed and stored in 500 µl of Tris, pH 8.0 as described above for the BirA*-KH BioID experiment.

Trypsin digestion, tandem mass tag labeling, and analysis of tryptic peptides by mass spectrometry is detailed in Supporting Information and a summary of the data is presented in Tables S2 and S3.

### Generation of Rabbit anti-KH antibody

A custom polyclonal anti-KH antibody was generated by GenScript, using their 49-day antibody generation protocol. Briefly, two rabbits were injected with 200 µg of His_6_-tagged recombinant KH fragment (rKH), representing amino acids 31-343, emulsified in Freund’s complete adjuvant. The rabbit was boosted 3 times at 14-day intervals with 200 µg rKH emulsified in Freund’s incomplete adjuvant. Antibody specificity for KH was evaluated by Western blot comparing the reactivities of the rabbit serum from immunized rabbits to *L. mexicana* cell lysates from wild-type and *∆kh* parasites (Fig 4A), normalized to tubulin levels detected using mouse anti-α-tubulin antibody 1:10,000, monoclonal B-5-1-2, (Sigma-Aldrich, catalog number T5168). Blots were probed with secondary antibodies goat anti-rabbit-HRP (Sigma, Cat#A0545) and goat anti-mouse-HRP (Jackson ImmunoResearch Laboratories, Cat#115-035-174) at 1:10,000 dilution. SuperSignal® West Pico Chemiluminescent Substrate (Thermo Fisher) was used for detection, and an Image Quant LAS 400 (GE Healthcare) scanner was employed to acquire luminescent images. Images were analyzed using ImageJ, version 2.0.0-rd-69/1.52p.

### HIS::KH pulldowns

Parasites expressing HIS::KH and Ty1::KHAP1, Ty1::KHAP2, or Ty1::23.2330 were grown to ~5 - 7 × 10^6^ cells/mL, and 1 - 3.5 × 10^8^ total cells were used in each experiment. Half the cells were formaldehyde crosslinked, and the other half treated in parallel as a control sample. The crosslinking, protein preparation, and HIS-cobalt resin incubation and washes were performed as described for the HBH::KH experiment above. Proteins bound to the His-cobalt resin were eluted by incubation in elution buffer with Imidazole (45 mM NaH_2_PO_4_, 8M Urea, 270 mM NaCl, 150 mM imidazole), and the eluate concentrated using Amicon Ultra-0.5 Centrifugal Filter Units, 10K NMWL (Millipore Sigma). Eluates were resolved by electrophoresis on a 4 – 12% bis-Tris gel (Invitrogen) after reversing the formaldehyde crosslinking by boiling the protein samples for 30 min prior loading. Then proteins were transferred onto a nitrocellulose membrane (Amershan) and immunodetected by using the following antibodies: Rabbit anti-KH, generated in this study, 1:5000; mouse anti-Ty1 1:500 (47). Blots were then probed with secondary antibody and developed as described above for the western to test the specificity of the rabbit anti-KH antibody.

### Proximity ligation assay

Promastigotes were grown to a density of 1 - 5 × 10^6^ and fixed and permeabilized on coverslips as for the immunofluorescence analysis described below. From these fixed cells the manufacturer’s Duolink PLA Immunofluorescence protocol was followed closely using the Duolink In Situ Red Starter Kit Mouse/Rabbit (Millipore Sigma). Briefly, after blocking cells were incubated with two primary antibodies as indicated, rabbit anti-KH 1:250 (this study), and mouse anti-Ty1, 1:250 (47). As a negative control, PLA was performed with only one primary antibody, anti-KH, 1:250. Then PLA species-specific secondary antibodies with minus and plus probes were added followed by ligation, amplification, and hybridization to fluorescent oligonucleotides to allow fluorescent detection. Following the PLA reaction cells were mounted and imaged as described for immunofluorescence analysis below.

### Immunofluorescence analysis

For all localization analysis on fixed promastigotes, parasites were grown to 1 - 5 × 10^6^ cells/mL, washed once in PBS, and then allowed to settle on poly-lysine coated coverslips. Attached parasites were washed twice in PBS, then fixed in 4% formaldehyde (Ultra Pure EM grade, Polysciences, Warrington, PA) diluted in PBS for 10 min at room temperature followed by a 5 min incubation with 250 mM glycine to stop the fixation reaction. Coverslips were washed with PBS, then cells were permeabilized with 0.1% Triton X-100 (Sigma) diluted in PBS. Cells were incubated in blocking solution (PBS plus 4% goat serum, 0.01% saponin) for 1h at room temperature and washed 3 times with PBS. Then cells were incubated with the indicated primary antibodies diluted in blocking solution for 1 h at room temperature as follows: rabbit anti-OLLAS, 1:250 (GenScript, catalog no. A01658-40), mouse anti-HA 1:500 (BioLegend, catalog no. MMS-101R), mouse anti-Ty1, 1:500 (47), rabbit anti-KH, 1:250 (this study), mouse anti-α-tubulin 1:1000 (Sigma-Aldrich, catalog no. T5168), mouse anti-β-tubulin clone KMX-1, 1:2000 (Millipore, catalog no. MAB3408). After three washes with PBS, cells were incubated with a 1:1000 dilution of secondary antibodies coupled to Alexa Fluor dyes (Molecular Probes) as follows: Alexa Fluor® 488 goat anti-mouse IgG (H+L) (catalog no. A11029), Alexa Fluor® 595 goat anti-mouse IgG (H+L) (catalog no. A11005), and Alexa Fluor® 594 goat anti-rabbit IgG (H+L) (Invitrogen catalog no. R37117), as indicated, in blocking solution for 1 h at room temperature in the dark. After three washes in PBS, coverslips were mounted onto slides using DAPI Fluoromount-G (SouthernBiotech). When parasites were expressing fluorescent fusion proteins, cells were not permeabilized and no antibody detection was used to visualize those proteins, since the protein fluorescence was well maintained following formaldehyde fixation.

For immunofluorescence of amastigotes, parasites were allowed to infect THP-1 macrophages in T-25 cell culture flasks and replicate for four days (see description of macrophage infections below). Infected macrophages were washed with cold PBS, 2 mM EDTA, then scraped off the flask surface in 2 ml cold PBS, 2 mM EDTA. Infected macrophages were passed through 27-gauge needle five times and then through a 30-gauge needle ten times, and macrophage lysis was confirmed by light microscopy. Cells were washed in cold PBS, pelleted at 1000 × g for 5 min, and the pellet was resuspended in 250 µl of PBS plus 2 mM EDTA. This cell suspension was layered onto a 250 µl 90% Percoll cushion (Sigma) and centrifuged at 21,000 × g for 10 min. Amastigotes were collected from the interface and fixed in 4% formaldehyde. Fixed amastigotes were adhered to poly-lysine coated coverslips, and the immunofluorescence analysis was performed as described for promastigotes.

### Live-cell microscopy

To assess protein localization in immobilized live cells promastigotes were grown to 1 − 5 × 10^6^, washed once in PBS, and resuspended in 1/50 the original volume. These concentrated cells were added to PBS-primed CyGEL (BioStatus) at 1:10 v/v dilution. The mixture was pipetted onto cold slides, flattened with a coverslip, allowed to spread, and then warmed to solidify the CyGEL. These slides were then imaged directly.

### Fluorescence microscopy

All fluorescence images were acquired on a high resolution wide field Core DV system (Applied Precision). This system is an Olympus IX71 inverted microscope with a proprietary XYZ stage enclosed in a controlled environment chamber; DIC transmitted light and a solid state module for fluorescence. The camera is a Nikon Coolsnap ES2 HQ. Each image was acquired as Z-stacks with a 60× 1.42 NA PlanApo lens. The images were deconvolved with the appropriate optical transfer function using an iterative algorithm of 10 iterations.

Pearson’s Correlation Coefficients (PCCs) were calculated using the Imaris Coloc module using deconvolved images from the Core DV system. The calculations were limited to regions-of-interest defined by the fluorescent signal of KHAP1/2 in the cells to exclude the non-cell background in the image which can artificially inflate the PCC (48). As a positive control the same primary antibody was probed with two different secondary antibodies and their PCC calculated. As a negative control one channel was rotated 90° and then the overlap between the two signals was calculated. The analysis was performed for >20 cells per condition in 5 images, and the complete results can be found in Figure S1.

### Cytoskeleton Extraction

1 × 10^7^ parasites of indicated genotypes were spun down at 1000 × g for 10 min, then washed once in PBS and resuspended in 100 µl 1% NP-40 in PEME buffer (0.1M PIPES, pH 6.9; 2 mM EGTA; 1 mM magnesium sulfate; 0.1 mM EDTA, plus protease inhibitors (Halt Protease Inhibitor cocktail, Thermo Fisher)) and incubated at room temperature for 5 minutes. Samples were then centrifuged at 200 × g for 10 min. The supernatant was saved and the pellet was washed twice in PBS and then resuspended in 100 µl RIPA buffer (150 mM NaCl, 25 mM Tris, 1% Nonidet P-40, 0.5% sodium deoxycholate, 0.1% SDS plus protease inhibitors). Equivalent volumes of the pellet and supernatant fractions were run for each sample and analyzed by Western blot probed with rabbit anti-KH, generated in this study, 1:5000; and mouse anti-α-tubulin 1:10,000 (monoclonal B-5-1-2, Sigma-Aldrich). To image the blot the secondary antibodies goat anti-rabbit IgG IRDye 800CW (LI-COR, catalog number 926-32211) and goat anti-mouse IgG IRDye 680RD (LI-COR, catalog number 926-68070) were used at 1:10,000 dilution. All antibodies were diluted in Aqua Block buffer (Abcam) containing 0.1% Tween® 20. The blots were scanned for infrared signal using the Odyssey Infrared Imaging System (LI-COR Biosciences).

### Macrophage Infections

The human acute leukemia monocyte cell line (THP-1) was cultivated in RPMI 1640 medium (Invitrogen) supplemented with 10% heat-inactivated FBS (Thermo Scientific Hyclone, Logan, UT), 25 mM HEPES, 1% L-glutamine, 50 mM glucose, 5 mM sodium pyruvate, and 1% streptomycin/penicillin at 37 °C and 5% CO_2_. The cultures were diluted every 3 days to prevent cell count from exceeding 1 × 10^6^ cells/ml. Cells were kept for a maximum of 10 subcultured dilution cycles. THP-1 cells were differentiated with 100 ng/ml of phorbol 12-myristate 13-acetate (Sigma) for 48 h at 37 °C, 5% CO_2_. Differentiated THP-1 cells are adherent and were seeded in 4-well Lab-TekII Chamber Slides (Nalgene Nunc International, Rochester, NY) at a density of 3 × 10^5^ cells per well. *L. mexicana* promastigotes at stationary phase were added to the plates (1:10 macrophage/parasite ratio) and incubated for 1 day, 4 days, and 7 days at 34.5 °C, 5% CO_2_. At each time point, slides were fixed with methanol and stained using the HEMA3 STAT PACK staining kit as described by the manufacturer (Fisher Scientific). Infected macrophages were examined using a Nikon Eclipse 50i microscope equipped with a ×100 1.25NA oil objective (Nikon Instruments, Melville, NY), and the number of parasites/100 macrophages were determined by counting 300 cells in each of the triplicate experiments per round of infection. Images were captured using a Leica DM4 B upright wide-field fluorescence microscope with a Leica DMC2900 color camera (3.1 Megapixel CMOS sensor) and a 100x oil objective, NA 1.25, using the LAS V4.12 software.

## The abbreviations used are

KH: KHARON
KHAP: KH Associated Protein
TAP: tandem affinity purification
HBH: hexa-histidine-biotinylation domain-hexa-histidine epitope tag
ORF: open reading frame
mNG: mNeon Green fluorescent protein
PCR: polymerase chain reaction
DAPI: 4′,6-diamidino-2-phenylindole
PLA: proximity ligation assay
HYG: hygromycin resistance gene
SAP: spindle associated protein
IFA: immunofluorescence analysis
FAZ: flagellum attachment zone
FM: flagellar membrane
PPM: pellicular plasma membrane

## Author Contributions

FDK, KT, MAS, SML conceptualization; FDK, KT, JH, KS, MAS, and SML data curation; FDK and SML writing the original draft; SML, project administration; FDK, MAS, and SML, editing.

## Acknowledgements

We acknowledge the Oregon Health & Science University Proteomics Shared Resource and its director Larry David for performing the proteomics experiments. We appreciate the expert advice and support of the staff of the Advanced Light Microscopy Core in the Jungers Center for Neurosciences at Oregon Health & Science University. We thank Philip Yates and Kat Ng for critical readings of the manuscript.

## Notes

This work was supported by National Institutes of Health grants R01AI121160 and R21AI107144 (to S.M.L.).

The authors declare that they have no conflicts of interest with the contents of this article. The content is solely the responsibility of the authors and does not necessarily represent the official views of the National Institutes of Health.

